# A legacy role for DNA binding of Lon protects against genotoxic stress

**DOI:** 10.1101/317677

**Authors:** Rilee D. Zeinert, Justyne L. Ogdahl, Jing Liu, Qiyuan Yang, Yunguang Du, Benjamin B. Barros, Peter L. Freddolino, Cole M. Haynes, Peter Chien

**Affiliations:** Department of Biochemistry and Molecular Biology, Molecular and Cellular Biology Program, University of Massachusetts Amherst, Amherst MA; Department of Molecular, Cell and Cancer Biology. University of Massachusetts Medical School, Worcester MA; Department of Biological Chemistry and Department of Computational Medicine & Bioinformatics, University of Michigan, Ann Arbor, MI 48109, USA

## Abstract

DNA binding proteins are essential for cellular life, but persistently bound complexes have toxic consequences. Here we show that the proteotoxic responsive bacterial protease Lon clears proteins from DNA to promote genotoxic stress resistance. Purified Lon binds DNA and degrades neighboring bound proteins, while a fully active DNA-blind Lon variant does not. This variant can degrade substrates as normal during unstressed growth, complements pleotropic phenotypes of Δ*lon*, including proteotoxic resilience, but remains sensitive to genotoxic stresses and fails to degrade proteins efficiently during DNA damage. Transposon sequencing reveals that Δ*lon* is vulnerable to loss of protein-DNA eviction factors and we use dynamic nucleoid occupancy profiling to show that chromosome-wide protein turnover relies on Lon DNA binding. Finally, disrupting Lon binding to mitochondria genomes also results in genotoxic stress sensitivity, consistent with the bacterial ancestry of this organelle. We propose that clearance of persistent proteins from DNA by Lon originated in free-living α-proteobacteria and maintained during the evolution of mitochondria.

**Summary:** DNA binding by the Lon protease protects against genotoxic damage in a manner preserved from bacteria to mitochondria.

## Introduction

Maintenance, replication, and readout of genomes requires DNA-binding proteins, which perform their functions with specific dynamics. However, proteins that dwell on DNA beyond their normal lifetimes can result in toxic consequences such as collisions of DNA polymerase with these roadblocks. Given that many proteins are known to be associated with the DNA, cells must have dedicated protein eviction pathways to prevent DNA-Protein roadblocks. In eukaryotes, chromatin remodeling factors evict nucleosomes and recent discoveries of the Sprtn proteases that remove covalently bound protein adducts from DNA represent the most extreme version of this process (Reinking et al., 2020). In bacteria, chromatin is condensed into the nucleoid which is highly occupied with both specific binding proteins (e.g., transcription factors) and less specific factors (e.g. HU, H-NS, etc). Like in eukaryotes, clearance of these proteins is needed, however, what factors collaborate to control this process in bacteria is less understood.

In bacteria, the AAA+ protease Lon degrades misfolded proteins (Goldberg, 1972; Gur and Sauer, 2008) and seminal work in *E. coli* revealed that loss of Lon sensitized cells to genotoxic stresses (Mizusawa and Gottesman, 1983; Witkin, 1946). Lon also plays a role during normal growth. For example, in the alpha-proteobacteria *Caulobacter crescentus (Caulobacter*), the Lon protease degrades the epigenetic regulator CcrM (Wright et al., 1996), the transcriptional regulator SciP (Gora et al., 2013), and the replication initiator DnaA (Jonas et al., 2013), in addition to its role in protein quality control (Lu et al., 2007). However, whether Lon plays a role in genotoxic tolerance in bacteria other than *E. coli* remains poorly understood. In mitochondria, which have an alphaproteobacteria origin (Gray, 2012), Lon has been clearly shown to manage the response to protein misfolding but it is unclear if it regulates mtDNA stress responses (Bota and Davies, 2002; Lu et al., 2007; Nargund et al., 2012). Intriguingly, early biochemical work with Lon shows that it can bind DNA (Charette et al., 1984; Chung and Goldberg, 1982; Liu et al., 2004; Lu et al., 2007) and more recent work identified specific residues important for DNA binding (Karlowicz et al., 2017), but the biological consequences of this binding remains unclear.

Here we report that Lon binding to DNA is important for genotoxic stress responses but not proteotoxic stress responses. Using transposon sequencing, we find that factors important for evicting persistent protein complexes from DNA show synthetic fitness defects with loss of Lon DNA binding. In vitro, purified Lon degrades neighboring proteins when bound to the same DNA region and genome-wide proteome occupancy shows that Lon binding to DNA affects the lifetime of proteins on the chromosome. Finally, we show the importance of Lon DNA binding in genotoxic stress tolerance is preserved in mitochondria, highlighting the link between these organelles and their a-proteobacteria ancestor. Overall, our work supports a model where Lon degrades lingering neighbor proteins when bound to chromosomal DNA, a process that is especially important during the response to genotoxic stresses.

## Results

### DNA binding of Lon is important for genotoxic stress tolerance

We generated strains expressing only a DNA-blind variant of Lon (Lon4E) by mutating residues implicated in DNA binding based on homology to the *E. coli* protein (Karlowicz et al., 2017). Cells lacking Lon exhibit profound morphological defects including filamentation, loss of motility in soft agar (Wright et al., 1996), and sensitivity to proteotoxic stresses such as canavanine (Zeinert et al., 2020) (Figure 1A). Expression of Lon4E largely rescues these phenotypes (Figure 1A,C; Figure S1). Unlike WT Lon, purified Lon4E fails to bind DNA (Figure 1B) but is completely active for degradation of model unstructured substrates such as casein (Figure S1). *In vivo*, loss of Lon results in increased levels of its substrates DnaA and CcrM (Jonas et al., 2013; Wright et al., 1996) and Lon4E restores steady-state levels to essentially wildtype levels (Figure S1).

**Figure 1.**
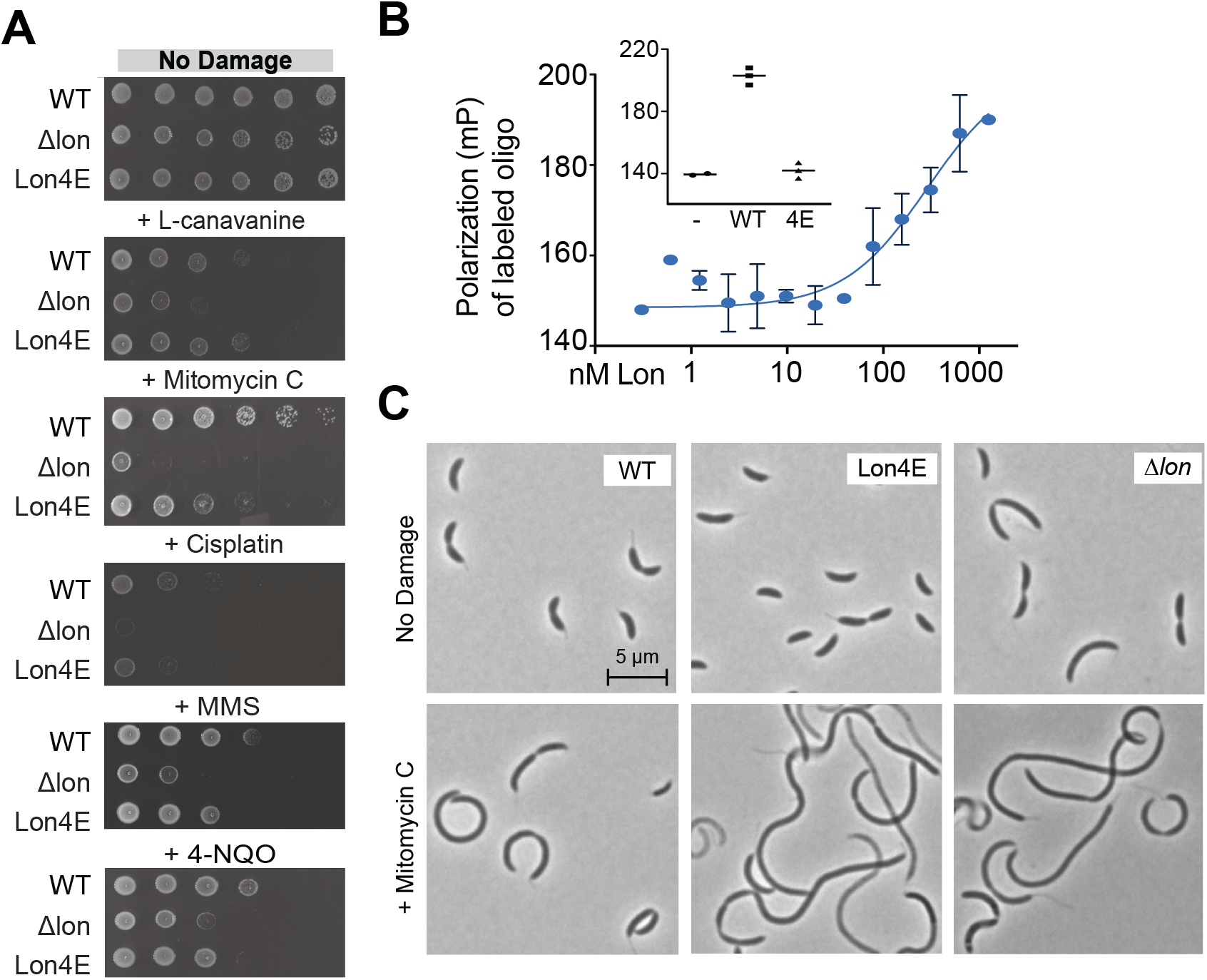
**A**. Serial dilutions (10-fold dilutions left to right) of wildtype (WT), Lon deficient (Δ*lon*), or DNA binding deficient Lon expressing (Lon4E) strains on media containing various concentrations of either proteotoxic (L-canavanine (L-can)) or genotoxic (mitomycin C (MMC), cisplatin (Cis), methyl methanosulfone (MMS), 4-nitroquinolone (4NQO)) compounds. **B.** Lon binding to fluorescently labeled DNA oligo as measured by change in fluorescence polarization. Data was fit to a hyperbolic binding model, K_D_= 286 nM (95% CI: 90-1200 nM). Concentration is reported as hexamers of Lon. inset: polarization measurement of labeled DNA in the presence of wildtype Lon or Lon4E. **C.** Morphology of cells during normal growth and after damaging with mitomycin C. Scalebar showing 5 μm in first panel, all other panels are the same magnification.

Interestingly, although they are resistant to proteotoxic stresses, Lon4E strains remain sensitive to DNA damaging stresses, most particularly genotoxic agents that yield complex damages such as mitomycin C (Figure 1A). Consistent with this, the prolific filamentation upon DNA damage associated with Δlon strains is also seen with Lon4E but not WT strains (Figure 1C). Our general interpretation is that the Lon4E allele is fully capable of complementing wildtype Lon activity during normal and proteotoxic stress but fails to fully protect against genotoxic stress.

### Loss of protein eviction factors is synthetically sick with loss of Lon activity

We had previously used transposon sequencing to identify genes that are differentially critical in a Lon-dependent manner (Figure 2A; (Zeinert et al., 2020)). Given the extreme sensitivity of Lon-deficient strains to genotoxic agents (Figure 1), we hypothesized that many members of the DNA damage response (DDR) pathway would be essential in Δ*lon*. However, we found that most genes in the DDR were unaffected, including the sentinel protein RecA, translesion polymerase (*imuB*) and mitomycin C specific responders (*mmcA*) (Figure 2 A,B), which we confirmed by generating viable double mutants (Figure 2 C).

**Figure 2.**
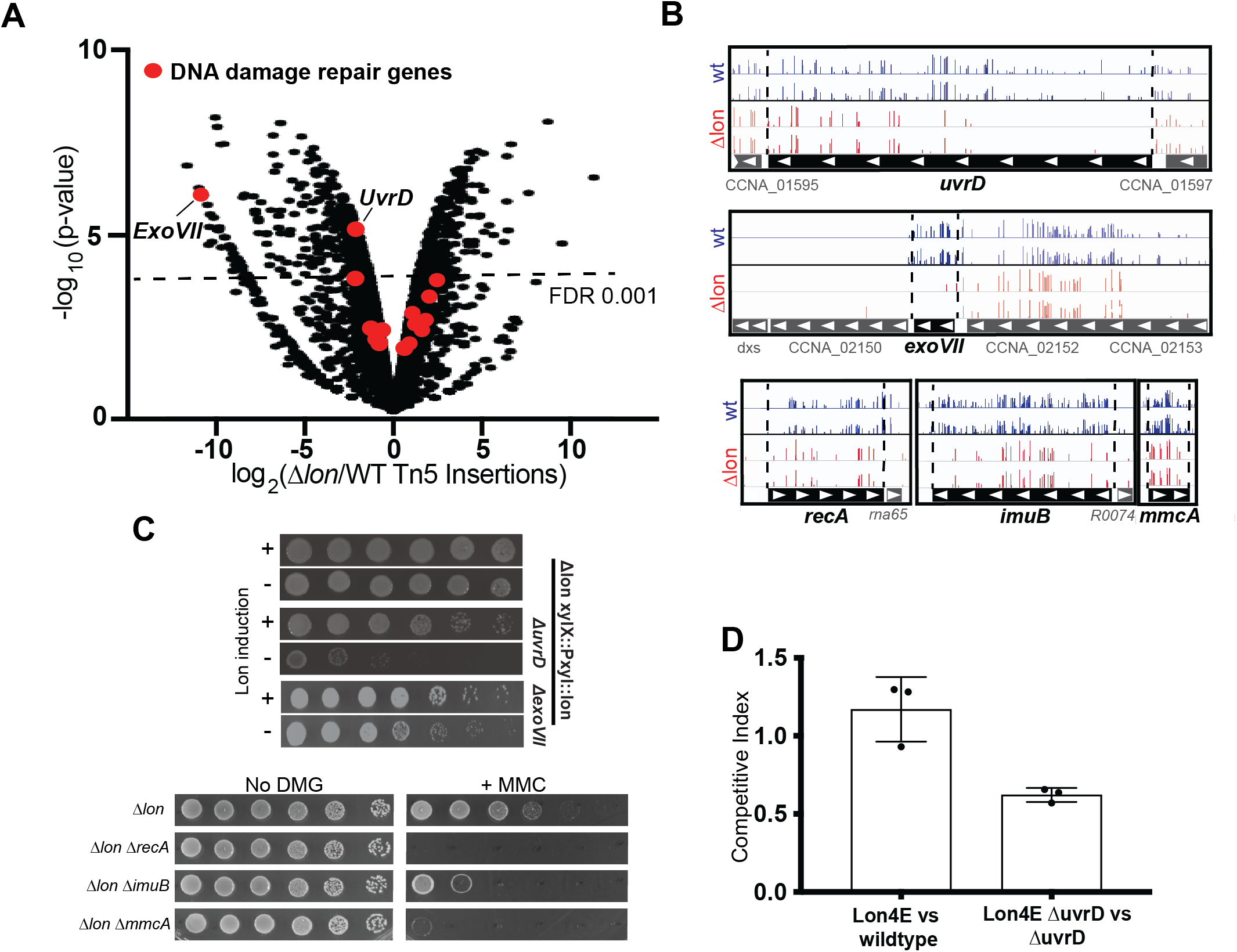
**A.** Transposon-sequencing reveals genes that differentially affect fitness in wildtype or Δlon strains (Zeinert et al., 2020), highlighting genes annotated as DNA damage repair genes (red). **B.** Snapshots of transposon insertion sites for DNA damage repair genes in either wildtype (WT) or Δlon strains. **C.** above: Induction (+) or depletion (-) of Lon in either wildtype, ΔuvrD, or ΔexoVII strains, showing synthetic sickness. below: Deletions of DNA damage repair genes *recA, imuB*, or *mmcA* can be generated in Δlon strains without loss of viability and as expected are generally important for response to MMC. D. Lon4E strains grow as well as wildtype based on competition, but deletion of *uvrD* reduces fitness in a Lon4E background.

Only two DDR genes passed our filter as more essential in a Δ*lon* background (Figure 2 A,B). The first of these is *uvrD*, a helicase originally known for its role in DNA damage repair (Ogawa et al., 1968). UvrD can evict proteins bound to DNA, such as disassembling RecA nucleofilaments (Petrova et al., 2015; Veaute et al., 2005) and stalled RNAP complexes (Epshtein et al., 2014). While we could delete *uvrD* from wildtype cells, we could not delete *uvrD* from Δ*lon* cells and depletion of Lon in a *DuvrD* strain was lethal (Figure 2C). Similarly, we found *xseB*, a subunit of exonuclease VII that is important for reactivation of stalled replication forks and mismatch repair (Lovett, 2011) also results in synthetic sickness when Lon is depleted (Figure 2C; Figure S2).

One pathway where UvrD and ExoVII converge is in clearing protein roadblocks from DNA (by UvrD) before they cause stalling of replication forks that must be restarted (initiated by XseB). Interestingly, while Lon4E strains were equally fit as wildtype strains under normal conditions, loss of UvrD in a Lon4E background compromised growth compared to the parental Δ*uvrD* strain (Figure 2D). Taken together, we hypothesize that Lon binding to DNA aids in the clearance of bound proteins, a function of particular importance during response to DNA damage. We therefore looked at how protein dynamics *in vivo* could be affected by Lon DNA binding.

### Lon binding to DNA affects condition dependent protein degradation

Degradation of known Lon substrates, such as DnaA and CcrM, during nondamaging conditions was the same in Lon4E strains as wildtype (Figure 3A). However, when MMC was added to induce DNA damage, wildtype cells maintained robust DnaA turnover, while in Lon4E, DnaA is degraded at ~50% of the wildtype rate. This stabilization did not extend to a different Lon substrate, CcrM (Figure 3A), nor was this effect seen with the ClpXP substrate CtrA (Jenal and Fuchs, 1998; Joshi et al., 2015) (Figure S3).

**Figure 3.**
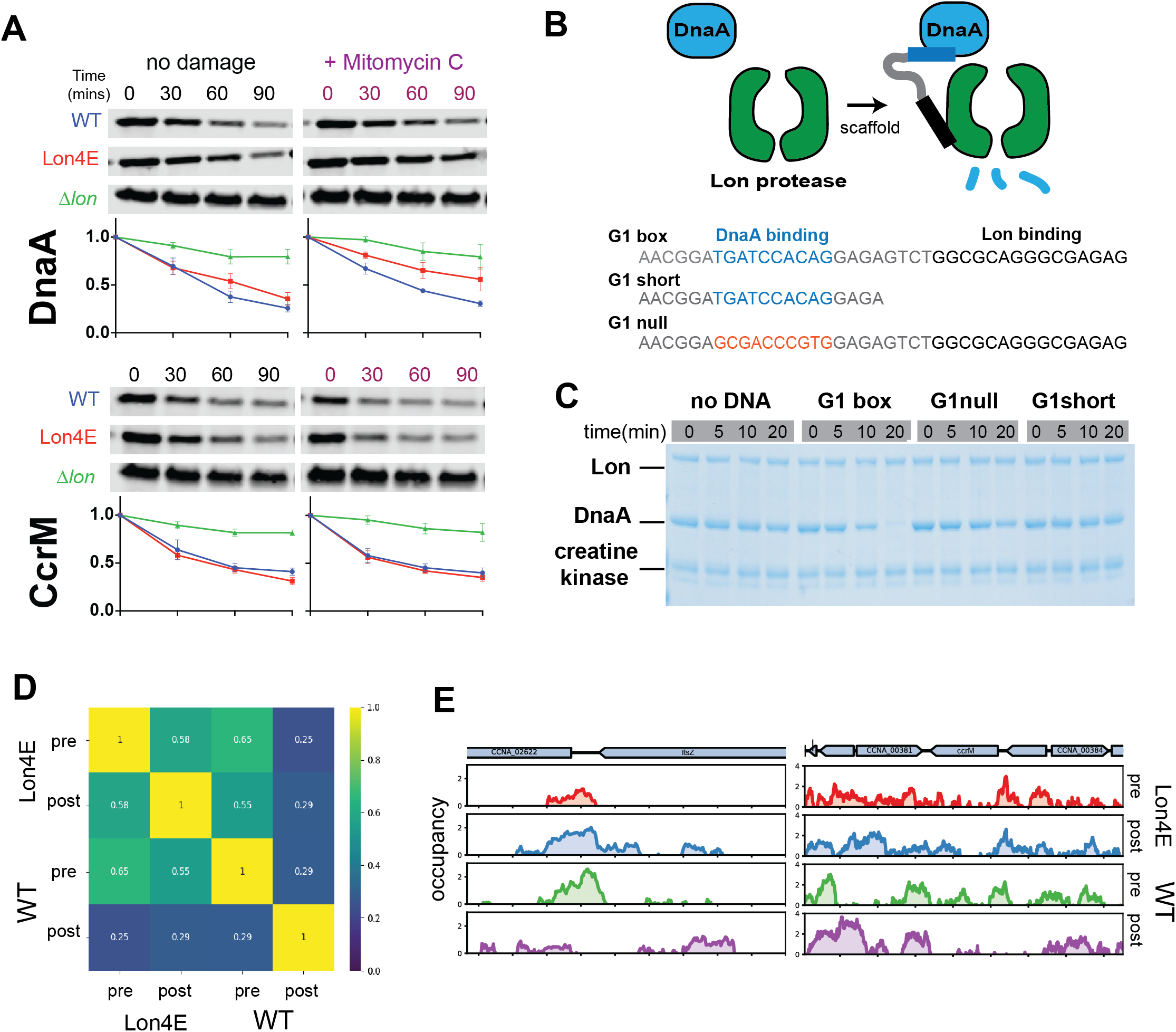
**A.** Degradation of DnaA and CcrM in wildtype, Lon4E or Δlon strains as monitored by translational shutoff experiments either in nondamaging or DNA damaging (+Mitomycin C) conditions. B. Cartoon illustrating dsDNA oligonucleotides used to test for scaffolding. C. Degradation of purified DnaA by Lon in the presence of different dsDNA scaffolds. D. Spearman correlation matrix of genome occupancy in wildtype (WT) or strains expressing Lon4E (Lone4E) as monitored by IPOD (Freddolino et al., 2021) either before (Pre) or two hours after (Post) translational shutoff. Degree of correlation is shown by color and inset values. E. Representative tracks showing changes in protein occupancy at *ftsZ* and *ccrM* neighboring regions in either WT or Lon4E strains.

*In vitro*, purified Lon degrades DnaA poorly on its own consistent with prior work showing that unfolded substrates activate Lon degradation of DnaA (Jonas et al., 2013). Interestingly, addition of a dsDNA with canonical DnaA binding sites results in robust degradation of DnaA by Lon (Figure 3C). This effect required both an intact DnaA binding motif and a sufficiently long adjacent region for Lon to bind as well (Figure 3C). Consistent with a requirement for DNA binding by Lon for this activity, purified Lon4E does not degrade DnaA as well as wildtype in the presence of this dsDNA scaffold (Figure S3). We conclude that Lon binding to DNA is needed for robust degradation of neighboring substrates, such as DnaA, an effect particularly important during genotoxic stress.

### Locus specific protein occupancy of the chromosome is regulated by DNA binding of Lon

Our working model is that DNA binding AAA+ proteases like Lon can clear persistent protein complexes. If this is true, then Lon mutants deficient in DNA binding should show persistent protein occupancy on the chromosome. To explore this, we used a modification of IPOD, a technique where fragmented crosslinked protein-DNA is purified and sequenced to reveal protein occupancy of the bacteria chromosome (Freddolino et al., 2021). We monitored chromosome occupancy in wildtype and Lon4E strains following translation inhibition, to reveal loci where DNA-bound proteins are removed by AAA+ proteases.

Our dynamic IPOD approach revealed many regions of the chromosome that become less occupied after translational inhibition in a wildtype strain - many of these regions remained occupied in a Lon4E strain after similar treatment (Figure 3D). Correlation analyses show that the occupancies of pre-inhibition wildtype, pre-inhibition Lon4E, and post-inhibition Lon4E were more like each other compared to the occupancy seen in the wildtype post-inhibition sample (Figure 3C). Closer examination of occupancy data suggests a role for Lon clearing gene regulatory regions, such as those present in the promoter regions of *ccrM* and CCNA_02622, a putative amidase located adjacent to *ftsZ* (Figure 3D). Taken together, these data suggest that regions of the genome are bound with proteins that are cleared by Lon when it binds to DNA.

### DNA binding properties of Lon are important for genotoxic stress responses in mitochondria

Mitochondria have an endosymbiotic origin (Margulis, 1970) and the current model is that a free-living alpha-proteobacterium was engulfed by the ancestor of the modern eukaryotic cell (Gray, 2012). Because *Caulobacter* is an a-proteobacteria, we explored if the need for DNA binding of Lon in tolerating DNA damage was preserved in mitochondria. In *C. elegans*, mitochondrial Lon (LONP-1) is required for development and growth (Nargund et al., 2012). Based on homology, we engineered a variant of *lonp-1* with mutations in the corresponding predicted DNA binding residues to generate LONP-1^4E^ and used CRISPR/Cas9 to produce worms expressing only this allele at the endogenous locus. Importantly, we found that worms expressing only the LONP-1^4E^ variant at endogenous levels were healthy, grew and developed normally (Figure 4A,D). Because LONP-1 has a known role in mitochondrial protein quality control by recognizing and degrading diverse proteins that incur oxidative damage (Bota and Davies, 2002), we infer that LONP-1^4E^ is capable of maintaining proteostasis during animal development. Pull-down experiments indicated that LONP-1 robustly binds mitochondrial genomes *in vivo* as previously suggested (Fu et al., 1997; Lu et al., 2007), but LONP-1^4E^ does not (Figure 4B). Therefore, DNA binding of mitochondria Lon is not essential for normal development or growth.

**Figure 4.**
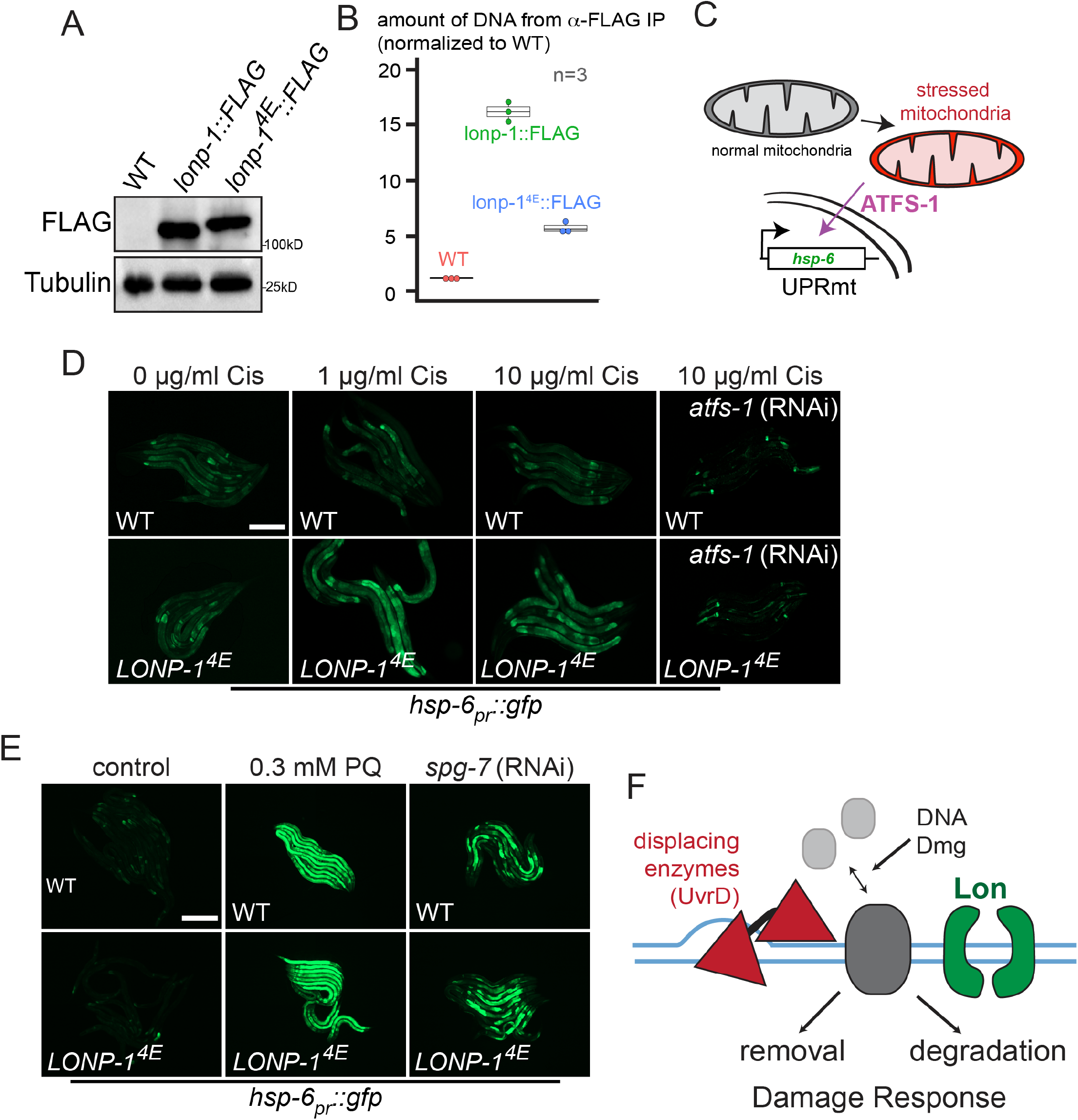
DNA binding mutants of mitochondria Lon. **A**. Immunoblots of lysates from WT worms, or strains expressing LONP-1-FLAG or LONP-1^4E^-FLAG. Tubulin serves as a loading control. **B**. ChIP of mtDNA with FLAG antibody in WT, LONP-1-FLAG and LONP-1^4E^-FLAG strains. **C.** Cartoon illustrating hsp-6 reporter of mitochondria stress. **D.** Photomicrographs of *hsp-6_pr_::gfp* WT or LONP-1^4E^ worms raised on DMSO or cisplatin, scale bar, 0.5 mm. **E.** Photomicrographs of *hsp-6_pr_::gfp* WT or LONP-1^4E^ worms raised on paraquat (PQ) or *spg-7*(RNAi). **F.** Model for how Lon binding to DNA clears persistent proteins (gray) together with other evicting enzymes such as UvrD (red).

The mitochondrial unfolded protein response (UPR^mt^) couples the state of mitochondrial protein homeostasis to an upregulation of nuclear encoded chaperone genes. Induction of the UPR^mt^ requires the transcription factor ATFS-1, which is targeted to the nucleus when mitochondria undergo stress. Complete depletion of LONP-1 activates the UPR^mt^, as determined by upregulation of the transcriptional mitochondrial chaperone reporter *hsp-6_pr_::gfp* (Yoneda et al., 2004) (Figure 4C) consistent with a role for LONP-1 in the maintenance of mitochondrial function (Nargund et al., 2012). LONP-1^4E^ worms do not activate the UPR^mt^ suggesting mitochondrial protein homeostasis is intact (Figure 4D). However, LONP-1^4E^ worms exposed to cisplatin showed a dramatic activation of UPR^mt^ that is ATFS-1 dependent while wildtype worms show no response (Figure 4D). Importantly, mitochondrial dysfunction driven by depletion of the protein quality control protease SPG-7 caused UPR^mt^ activation in both wildtype and LONP-1^4E^ animals to the same degree (Figure 4E), as does exposure to paraquat which induces mitochondrial oxidative stress (Figure 4E). These results show that, similar to what is seen in *Caulobacter*, loss of DNA binding of Lon in mitochondria results in a specific intolerance to genotoxic stress.

## Discussion

The role of DNA binding in Lon has been enigmatic since its initial observation (Chung and Goldberg, 1982; Zehnbauer et al., 1981). Our working model is that Lon binding to DNA allows it to recognize neighboring proteins and serves as an additional layer of quality control to reduce the residence time of persistent DNA binding complexes (Figure 4F). This oversight works in concert with other protein eviction factors such as UvrD and seems particularly important during DNA damaging conditions (Figure 4F). Using a genome wide approach, we find evidence for clearance of protein complexes during normal growth that relies on Lon binding to DNA. Finally, we have shown that the same properties of DNA binding in Lon that are important for genotoxic stress responses in *Caulobacter* are also important in mitochondria, consistent with the endosymbiotic α-proteobacteria ancestry of this organelle.

Overall, our working model is that Lon plays a canonical role in proteotoxic stress response by degrading misfolded or damaged proteins. This function occurs regardless of DNA binding. However, DNA binding of Lon also allows for local control of neighboring proteins that is particularly important during genotoxic stress conditions. Our working model is that Lon clearance of persistently bound proteins prevents the accumulation of protein roadblocks that limit replication or transcription. This is particularly important in bacteria and bacterially derived organelles, as replication, transcription and translation occur concurrently in the same compartment. Because Lon recognizes substrates based on general chemical properties as opposed to highly specific sequence motifs, Lon would be ideally suited to serve in this capacity as the identity of the protein roadblock will likely depend on the specific circumstance that generated it.

## Acknowledgements

We thank Lucy Shapiro for supplying the anti-CcrM antibody and members of the Chien lab and the Protein Homeostasis theme of the Center for Models to Medicine for helpful feedback.

## Author contributions

P.C., J.L., and R.Z. conceptualized the study. B.B., J.L., J.O., R.Z, Q.Y., and Y.D. performed experiments. P.C., C.H., R.Z, and J.L wrote the initial draft. All authors reviewed the final draft.

## Data and materials availability

All data is available in the main text or the supplementary materials.

## Materials and Methods

### Bacteria strains and media

*C. crescentus* was grown on peptone yeast extract (PYE) at 30°C. 1.5% agar was used for solid media. Carbon sources were supplemented at final concentrations of 0.2% glucose or xylose. Antibiotics were used at the following concentrations for solid media growth: Kanamycin 25 μg/ml, Spectinomycin 100 μg/ml, Tetracycline 1.5 μg/ml and Gentamycin 25 μg/ml. For growth of *E. coli* in solid or liquid media, we used lysogeny broth (LB) at 37°C. Antibiotics were used at the following concentrations: Chloramphenicol 30 μg/ml, Kanamycin 50 μg/ml, Tetracycline 15 μg/ml and Gentamycin 50 μg/ml. After initial selection steps for strain construction, antibiotic selection was excluded from all physiological related studies.

### Strain construction

Deletion strains were constructed using the two-step recombination procedure with counter selection by sucrose/*sacB*, using pNPTS138 plasmids containing deletion cassettes comprised of 1000 bp upstream and downstream of the coding regions flanking a gentamycin resistance cassette. Following transformation, initial primary selection was on PYE with Kanamycin 25 μg/ml. Primary colonies were grown overnight in PYE without selection to allow for secondary recombination. Overnight cultures were plated onto solid media supplemented with 3% (w/v) sucrose and 25 μg/ml Gentamycin. To validate gene deletion, resulting strains were tested for antibiotic sensitivity to Kanamycin and resistance to Gentamycin. Phage transduction of marked alleles was performed using PhiCr30.

Allelic replacement of Lon4E was performed by transforming pNPTS138-Lon4E into CPC546 (lon::specR) with primary selection on PYE with Kanamycin 25 μg/ml and secondary selection on PYE + 3% (w/v) sucrose. The resulting strains were genotyped for antibiotic sensitivity to Kanamycin and Spectinomycin. Clones harboring the correct genotype were then confirmed by PCR amplification of the *lon* locus, Sanger sequencing, and whole genome resequencing of the strain.

### Protein Purification and *in vitro* degradation assays

Lon, Lon4E, and DnaA were purified with protocols as previously described (Jonas et al., 2013) using hydroxyapatite resin (Sigma) instead of P11 cellulose for Lon/Lon4E.

FITC casein degradation was performed as previously described (Vieux et al., 2013). Briefly, degradation assays (30°C) were as follows unless noted otherwise: Lon/Lon4E hexamer 0.15 μM, 10 μg/ml FITC casein type II (Sigma), ATP 4 mM, 75 μg/ml creatine kinase, 5mM creatine phosphate. Final concentrations of components in degradation assays (30°C) were as follows unless noted otherwise: Lon/Lon4E 0.15 μM (hexamer concentration), DnaA 2.5 μM, DNA 2 μM, ATP 4 mM, 75 μg/ml creatine kinase, 5 mM creatine phosphate.

### Fluorescent polarization studies

ssDNA (AACGGATGATCCACAGGAGAGTCTGGCGCAGGGCGAGAG) labeled with fluorescein (FAM) was ordered from Integrated DNA Technologies. Wildtype Lon was titrated in two-fold dilutions starting from 1.25 μM (all concentrations in hexamer). For inset in Figure 1, 300 nM of Lon or Lon4E hexamer was incubated with 200nM FAM-DNA in 20mM HEPES, 10mM MgCl_2_, 100mM KCl with 0.05% Tween. Polarization measurements were read from 40 uL of these mixtures using opaque non-binding black 384-well plates (Corning) and a SpectraMax M5 plate reader (Molecular Devices), with excitation and emission wavelengths set at 460 nm and 540 nm respectively.

### Bacteria Characterization: Motility, Morphology, Viability and Stress Assays

For motility assays, single colonies were picked from PYE agar and inoculated into 0.3% PYE agar. Plates were grown at 30°C for 2 days before imaging.

Microscopy images are phase contrast images of exponentially growing *Caulobacter* cells OD_600_ 0.4 (Zeiss AXIO ScopeA1) mounted on 1% PYE agar pads using a 100X objective.

Caulobacter strains were cultured in PYE medium overnight. Cultures were diluted to OD_600_ 0.05 and grown to OD_600_ ≈ 0.4, after which 10-fold serial dilutions were spotted onto regular PYE agar or agarose (5-Azacytidine) plates containing the indicated concentrations of Mitomycin C, Cisplatin, L-canavanine, Methyl methanosulfonate, and 4-Nitroquinoline N-oxide (Sigma) followed by incubation at 30°C for 72 hours prior to imaging. Mitomycin C, L-canavanine, and Cisplatin was prepared at a stock concentration of 0.4 mg/ml, 100 mg/ml, and 20 mM respectively in water followed by filter sterilization. 4-NQO was prepared at a saturated solution in acetonitrile.

For synthetic lethality experiments with Lon depletion, xylose was present in all cultures at a final concentration of 0.2%. Overnight cultures were back diluted to OD_600_ 0.05 and outgrown to OD_600_ ≈ 0.4. Cells were 10-fold serially diluted, plated onto media containing 0.2% glucose or 0.2 xylose, grown for 72 hours at 30°C and imaged.

### Western Blot Analysis

Strains were cultured in PYE medium overnight. Cultures were diluted to OD_600_ 0.05 and grown to OD_600_ ≈ 0.4. For steady state or translation shutoffs (30 μg/ml chloramphenicol) cells were normalized and loaded onto a 10% BIS Tris pH 6.8 gel. Samples were run at 150V for 60 mins. Gels were transferred to nitrocellulose membranes at 100V for 50 minutes. Following transfer, blots were blocked with TBST with 5% milk for 45 minutes and probed with anti-rabbit primary CcrM, DnaA, and ClpP at 1:5,000 dilution overnight at 4°C. Blots were washed three times for 5 minutes in TBST. Followed by incubation with Goat anti-rabbit LiCOR 800CW secondary for 1 hour. Blots were washed three times in TBST and imaged using a LiCOR Odyssey Imager, quantification was performed using ImageJ (NIH).

### Worm strains and bacterial food sources

Hermaphrodite worms were raised on the OP50 strain of *E. coli* unless treated with *atfs-1*(RNAi), in which the HT115 *E. coli* strain harboring the *atfs-1*(RNAi) plasmid was used (Haynes et al., 2010). The UPR^mt^ reporter worm strain harboring the *hsp-6pr::gfp* transgene used in the *C. elegans* imaging experiments was described previously (Yoneda et al., 2004). All new strains were confirmed by sequencing, and backcrossed three times prior to crossing into the *hsp-6pr::gfp* background.

### *C. elegans* cisplatin experiments

Worm strains were synchronized by bleaching. The eggs were transferred to plates containing the described concentration of cisplatin (Sigma), allowed to hatch and develop for 36 hours, when they were harvested and imaged as described (Nargund et al., 2012).

### CRISPR/Cas9 genome editing

Via CRISPR-Cas9, a nucleic acid sequence encoding the amino acids - GS<u>DYKDDDDK</u>GS<u>DYKDDDDK</u>GS<u>DYKDDDDK</u>-, which yields 3xFLAG epitope sites (underlined) interspersed by -GS- linkers, was inserted at the 3’ end of the endogenous *lonp-1* open reading frame at the native locus as described (Friedland et al., 2013; Katic et al., 2015; Paix et al., 2017). crRNAs and donor templates were generated and purchased from Integrated DNA Technologies.

#### LONP-1::FLAG

##### crRNA sequence

/AlTR1/rArArGrArArArGrGrGrArCrArCrGrUrArUrUrArUrGrUrUrUrUrArGrArGrCrUrArUrGrCrU/AlTR2/

##### Donor template

ATATTCGATTCGTTTCGCATTATGATGAGCTCTACGAGCATCTCTTCCAAGGATCCGACTACAAGGACGATGACGACAAGGGATCTGACTACAAGGACGATGACGACAAGGGATCTGACTACAAGGACGATGACGACAAGTAATAATACGTGTCCCTTTCTTTCACCCCCAGTTAATACCTTTCAGATAGATTTTG

#### LONP-1^4E^

To generate LONP-1^4E^

Amino acid residues 467 (Arginine), 468 (Arginine), 473 (Lysine) and 474 (Lysine) were all altered to aspartic acid residues via CRISPR-Cas9 as described above.

##### crRNA sequence

/AlTR1/rArArCrUrGrArArArGrArCrGrArCrGrArUrUrCrUrGrUrUrUrUrArGrArGrCrUrArUrGrCrU/AlTR2/

##### Donor template

CTACCTGGAATGGTTGACATCAGTACCCTGGGGACTTACATCACCCGAGAATGAGGAACTTTCAGTTGCAGAAGAGGCTCTTGATGAAGGACATTATGGA

### *C. elegans* western blots and mtDNA ChIP

Western blots and ChIP experiments were performed as described (*51*), with several modifications. Synchronized worms were cultured in liquid and harvested at the L4 stage by sucrose flotation. Worms were resuspended in cold PBS + protease inhibitors (Roche) and homogenized. The homogenate was treated with 1.85% formaldehyde for 15 min to cross-link DNA and protein. Glycine was then added to a final concentration of 125mM for 5 min at room temperature to quench the formaldehyde. The pellets were washed twice in cold PBS + protease inhibitors, centrifuged at 2500 x G. Samples were sonicated using a Bioruptor (Diagenode) for 15 min at 4°C on high intensity (30s on, 30s off). Samples were transferred to microfuge tubes and spun at 13,000 x G for 15 min at 4°C. The supernatants were precleared with pre-blocked ChIP-grade Pierce™ magnetic protein A/G beads (Thermo Scientific) and then incubated with Monoclonal ANTI-FLAG^®^ M2 antibody (Sigma, F1804) or Mouse mAb IgG1 Isotype Control (Cell Signaling Technology, G3A1) overnight at 4°C. The antibody-chromatin complex was then concentrated with pre-blocked ChIP-grade Pierce™ magnetic protein A/G beads. After washing, DNA-protein crosslinks were reversed by incubation 65°C overnight followed by treatment with RNaseA at 37°C for 2 hours, and then proteinase K at 55°C for 2 hours. Finally, immunoprecipitated and input DNA were purified with ChIP DNA Clean & Concentrator (Zymo Research, D5205) and used as templates for qPCR. Relative enrichment folds of detected regions were determined after normalization to input and total mtDNA primers as described (Nargund et al., 2015).

qPCR primers to quantify *C. elegans* mtDNA following ChIP:

5’-GCAGTCTTAGCGTGAGGACATTA -3’

5’- AAACATAAAACATAAATAGAACTAACCA-3’;

### Dynamic IPOD

*In vivo* protein occupancy (IPOD) was measured as described (Freddolino et al., 2021) in wildtype or Lon4E *Caulobacter crescentus* at initial and two hours after addition of chloramphenicol to shut off translation. For Spearman correlation, occupancy scores were averaged across overlapping 50 kb intervals across the genome and comparisons made between timepoints and strains as shown. Wig files for occupancy data are provided in Supplemental information.

## SUPPLEMENTAL INFORMATION

### Supplemental Figure Legends

**Figure S1.**
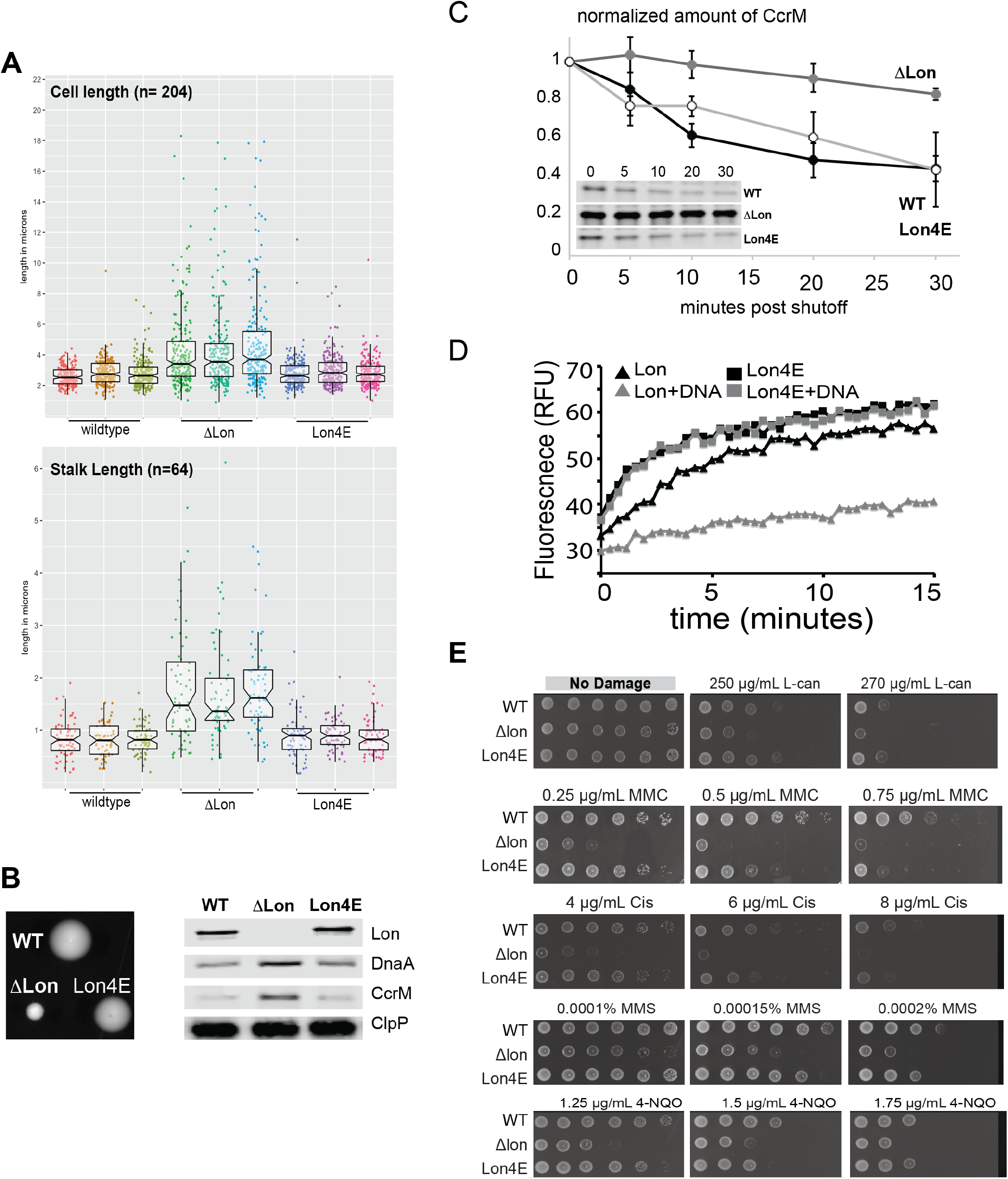
Supporting main Figure 1. **A.** Three biological replicates of wildtype, Δlon and Lon4E strains were individually grown to exponential phase. Cell length and stalk were measured for each. Cell lengths were determined automatically using MicrobeJ, Stalk lengths were measured manually using ImageJ (NIH). Individual measurements are shown as well as a notched box plot for these values. **B.** Growth in 0.3% Agar PYE media of WT, Δlon and Lon4E strains on left. Western blot of labeled proteins shown on right. **C.** Degradation of CcrM is restored in Lon4E. Chloramphenicol was added to exponentially growing cells and timepoints were taken at timepoints indicated. Quantification was performed as described in methods (n=3). **D.** Purified Lon or Lon4E can degrade fluorescently labeled casein (FITC-casein; Sigma) with similar efficiency. Degradation of FITC-casein results in increased fluorescence due to loss of self-quenching. Lon binding to certain single-stranded DNA oligonucleotides suppresses degradation of FITC-casein, but Lon4E is not responsive. **E.** Additional concentrations and replicates of growth under stress similar to main Figure 1.

**Figure S2.**
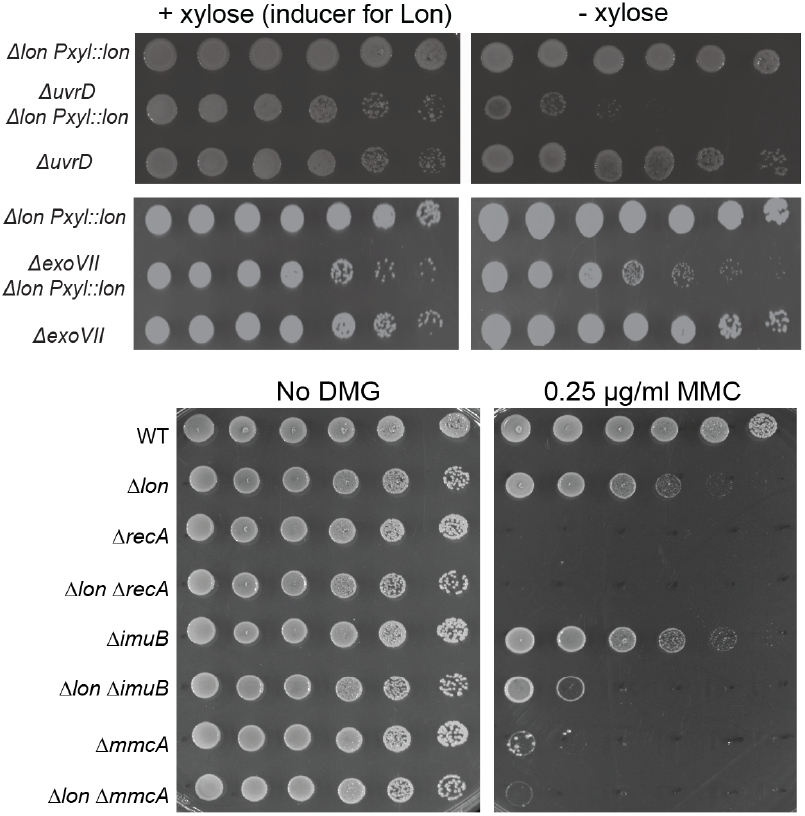
Supporting main Figure 2. Cells expressing a xylose inducible Lon as the sole copy (Δ*lon Pxyl:lon*) can tolerate loss of *uvrD* under inducing, but not repressing conditions. Similarly, depletion of Lon in a Δ*exoVII* strain also results in reduced fitness. Loss of Lon shows no synthetic effects with loss of *recA, imuB* or *mmcA*, although loss of these genes dramatically increases sensitivity to DNA damage produced by mitomycin C.

**Figure S3.**
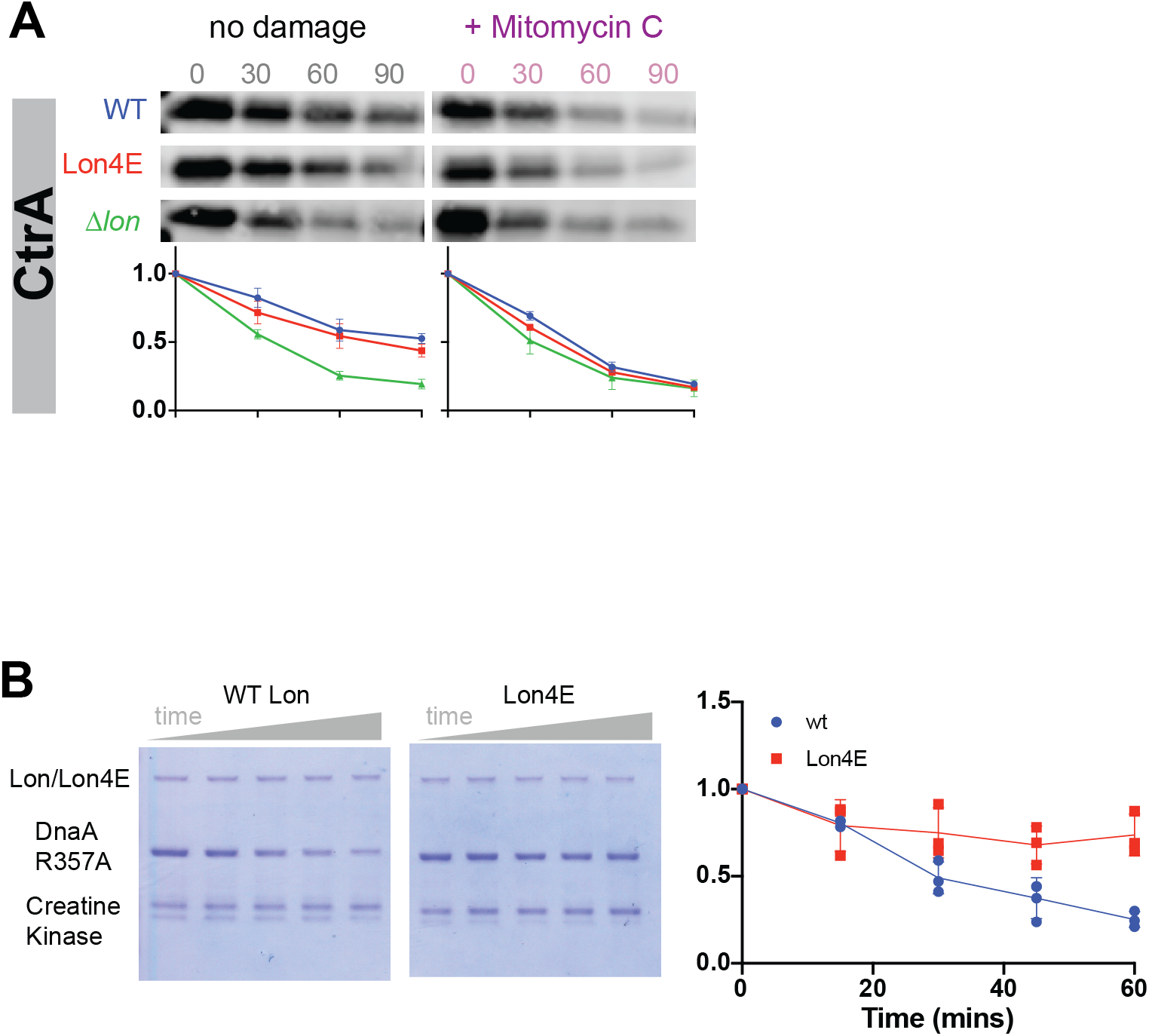
Supporting main Figure 3. A. CtrA degradation in WT, Lon4E and Δlon strains with and without mitomycin C damage. B. Degradation of DnaA by dsDNA is increased with WT Lon, but not with Lon4E. n.b., DnaAR357A is used here as it is unable to hydrolyze ATP and therefore is persistently locked in the ATP-bound / DNA-binding state.

